# Architectures of lipid transport systems for the bacterial outer membrane

**DOI:** 10.1101/064360

**Authors:** Damian C. Ekiert, Gira Bhabha, Garrett Greenan, Sergey Ovchinnikov, Jeffery S. Cox, Ronald D. Vale

**Author notes:** These authors contributed equally to this work (Damian C. Ekiert, Gira Bhabha). Correspondence (D.C.E.).

## Abstract

How phospholipids are trafficked between the bacterial inner and outer membranes through the intervening hydrophilic space of the periplasm is not known. Here we report that members of the mammalian cell entry (MCE) protein family form structurally diverse hexameric rings and barrels with a central channel capable of mediating lipid transport. The *E. coli* MCE protein, MlaD, forms a ring as part of a larger ABC transporter complex in the inner membrane, and employs a soluble lipid-binding protein to ferry lipids between MlaD and an outer membrane protein complex. In contrast, EM structures of two other *E. coli* MCE proteins show that YebT forms an elongated tube consisting of seven stacked MCE rings, and PqiB adopts a syringe-like architecture. Both YebT and PqiB create channels of sufficient length to span the entire periplasmic space. This work reveals diverse architectures of highly conserved protein-based channels implicated in the transport of lipids between the inner and outer membranes of bacteria and some eukaryotic organelles.

**HIGHLIGHTS:**

1. MCE proteins adopt diverse architectures for transporting lipids across the bacterial periplasm
2. Cryo-EM and X-ray structures reveal how the MlaFEDB complex, along with MlaC, might shuttle lipids across the periplasm
3. 3.9 Å cryo-EM structure of PqiB reveals a syringe-like architecture with a continuous central channel
4. YebT forms a a segmented tube-like structure, and YebT and PqiB are poised to directly link the inner and outer membranes to facilitate lipid transport.

## INTRODUCTION

The bacterial outer membrane (OM) serves a critical role as an environmental barrier, restricting the traffic of small molecules such as antibiotics into the cell (Nakae, 1976; Nikaido, 2003). Mutations that disrupt OM integrity reduce virulence in pathogenic bacterial species (Cox et al., 1999; Kong et al., 2012; Somerville et al., 1999; Wang et al., 2007) and increase their susceptibility to antibiotics and detergents (Helander et al., 1989; Kropinski et al., 1978; Nichols et al., 2011; Schnaitman and Klena, 1993), suggesting that targeting pathways important for OM maintenance and biogenesis may be a fruitful approach for development of new therapies for bacterial disease.

The OM of most gram-negative bacteria is asymmetric, with an outer leaflet rich in lipopolysaccharide (LPS) and an inner leaflet composed primarily of phospholipids (Kamio and Nikaido, 1976; Mühlradt and Golecki, 1975; Smit et al., 1975). Decades of research have uncovered a detailed, but still evolving, picture of the pathways and mechanisms involved in the biosynthesis, transport, and insertion of LPS into the OM (May et al., 2015; Ruiz et al., 2009). Yet, the mechanisms and structural basis of how nascent phospholipids are trafficked between the inner membrane (IM) and OM, and how asymmetry between the leaflets of the outer membrane is maintained remain unclear.

Proteins of the MCE superfamily [originally thought to mediate <u>M</u>ammalian <u>C</u>ell <u>E</u>ntry in *M. tuberculosis* (Arruda et al., 1993)] are defined by the presence of one or more conserved MCE domains at the sequence level, but have no similarity to other protein families of known structure or function. MCE proteins are ubiquitous among double-membraned bacteria (Casali and Riley, 2007) (Figure 1A) and are additionally found in eukaryotic chloroplasts (Awai et al., 2006), a bacteria-derived organelle surrounded by a double membrane. In contrast, MCE proteins are absent in bacteria bounded by a single membrane. MCE proteins are important virulence factors in TB and other bacterial pathogens (Carpenter et al., 2014; Gioffré et al., 2005; Joshi et al., 2006; Nakamura et al., 2011; Pandey and Sassetti, 2008; Rengarajan et al., 2005; Senaratne et al., 2008; Zhang et al., 2012), and have been suggested to play a role in the transport of lipids (Awai et al., 2006; Malinverni and Silhavy, 2009; Roston et al., 2011; Sutterlin et al., 2016; Xu et al., 2003, 2008), cholesterol and steroids (Klepp et al., 2012; Mohn et al., 2008; Pandey and Sassetti, 2008), and other hydrophobic molecules (Kim et al., 1998). An MCE protein from plants, and more recently from *E. coli*, has been shown to bind phospholipids (Awai et al., 2006; Thong et al., 2016). MCE systems are generally thought to drive the retrograde transport of hydrophobic molecules from the OM to the IM, although it has also been suggested that they may function as exporters to move molecules towards the outer membrane and the outside of the cell (Kim et al., 1998). As the direction of transport has only been inferred indirectly, it remains to be rigorously established whether MCE proteins mediate import or export, or whether this might vary between different MCE systems. Thus, MCE proteins appear to be critically important for OM function, but their precise role and mechanism of action remains unclear.

**Figure 1.**
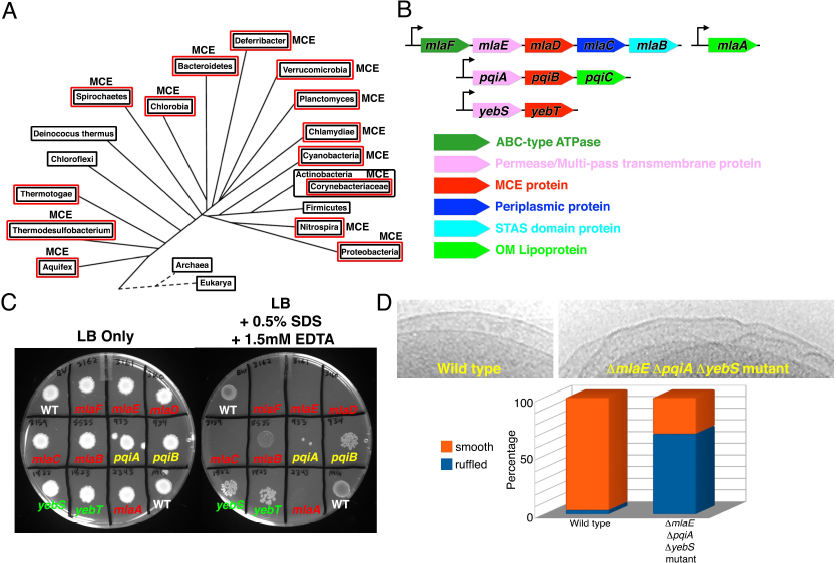
MCE proteins are important for outer membrane integrity. (A) Distribution of Pfam MCE proteins across bacterial lineages coincides with the presence of species with an OM (red boxes). (B) Organization of the *mlaFEDCB* and *mlaA*, *pqiABC*, and *yebST* operons from *E. coli*. (C) Mutation in the *mla* (red), *pqi* (yellow), and *yeb* (green) genes increase sensitivity to 0.5 % SDS + 1.5 mM EDTA, suggestive of an OM defect. (D) MCE mutants exhibit OM ruffling at a much higher frequency than wild-type cells (n∼50 WT and 50 mutant cells).

Here we report the X-ray and EM structures for the three MCE proteins from *E. coli*. One MCE protein, MlaD, forms a homo-hexameric ring, is part of a much larger multi-protein complex in the IM, and interacts with a soluble lipid-carrying protein, which likely serves as an intermediate that shuttles lipids between transport complexes in the IM and OM. In contrast, MCE proteins YebT and PqiB form two different, more elongated structures by stacking hexameric rings together to form barrels and tubes, which result in continuous channels that are of sufficient length to directly span the distances between the inner and outer membranes, without the need for a periplasmic shuttle protein. The distinct architectures of these MCE proteins suggest multiple strategies for moving lipids through the hydrophilic environment of the periplasm.

## RESULTS

### Mutations in MCE operons result in OM defects

The *E. coli* reference genome (strain MG1655) contains 3 MCE genes (Figure 1B): *mlaD* (<u>m</u>aintenance of OM <u>l</u>ipid <u>a</u>symmetry D), *pqiB* (<u>p</u>ara<u>q</u>uat <u>i</u>nducible B), and *yebT* (gene of unknown function; also known as MAM7). These MCE genes are part of three different operons, each of which also encode a multi-pass transmembrane protein (MlaE, PqiA, and YebS)(Figure 1B). The *mla* operon in addition encodes an ABC-type (ATP binding cassette) ATPase (MlaF), a predicted periplasmic protein MlaC, and a STAS domain containing protein, MlaB. The presence of an ATPase and a multi-pass transmembrane protein in some MCE operons has led to the hypothesis that MCE systems may encode a family of ABC transporters (Carpenter et al., 2014; Casali and Riley, 2007; Zhang et al., 2012). The *mlaFEDCB* genes function together with an OM lipoprotein, MlaA, encoded elsewhere in the genome (Figure 1B)(Malinverni and Silhavy, 2009). Recently, it has been shown that MlaFEDB interact to form a complex (Thong et al., 2016).

Mutations in the *mla* genes result in increased sensitivity to a combination of SDS and EDTA (SDS+EDTA), suggestive of a mild OM defect, which has been attributed to a loss of OM asymmetry (Malinverni and Silhavy, 2009). To assess whether other MCE proteins are similarly involved in OM maintenance or biogenesis, we constructed *E. coli* strains bearing single deletions of the MCE-encoding genes *pqiB* and *yebT*, or their associated putative permease genes, *pqiA* and *yebS*. These mutations in the *pqi* and *yeb* operons increased sensitivity of bacteria to 0.5% SDS + 1.25 mM EDTA (Figure 1C). We also constructed a triple mutant lacking the putative permease subunit from each operon (Δ*mlaE ΔpqiA ΔyebS*). The triple permease mutant is viable and shows no discernible growth defect relative to wild-type in liquid LB medium (Figure S1A). Thus, the Mla, Pqi, and Yeb systems are non-essential for *E. coli* growth in culture despite being critical for resistance to environmental stresses, including those likely encountered by pathogens during infection (Carpenter et al., 2014; Gioffré et al., 2005; Joshi et al., 2006; Nakamura et al., 2011; Pandey and Sassetti, 2008; Rengarajan et al., 2005; Senaratne et al., 2008; Zhang et al., 2012). The *ΔmlaE ΔpqiA ΔyebS* mutant exhibited SDS+EDTA sensitivity similar to the *ΔmlaE* single mutant. But in contrast to any of the single mutants, the triple mutant is more susceptible to outer membrane stressors, as this strain is uniquely sensitive to 5% SDS alone and, unexpectedly, to 100 mM CaCl_2_ (Figure S1B and S1C). As divalent cations are important for the stabilization of interactions between LPS molecules in the OM (Lugtenberg and Van Alphen, 1983), and the ratio of LPS to divalents may be important to maintain the appropriate degree of membrane fluidity/rigidity, and the mutant may be more sensitive to CaCl_2_ due to an already weakened OM.

Interestingly, exposure of *ΔmlaE ΔpqiA ΔyebS* mutant cells to the DNA stain DAPI resulted in strong nucleoid staining, whereas staining of wild-type cells was much weaker and largely confined to the cell periphery (Figure S1D). This result suggests that the function of the OM as a barrier to exclude hydrophobic molecules is severely compromised in the mutants. This is consistent with previous work suggesting that *E. coli* MCE mutants are more susceptible to hydrophobic antibiotics (Nichols et al., 2011). Staining of live cells with propidium iodide, a membrane-impermeable DNA dye which stains cells with a disrupted plasma membrane, confirmed that only a small population of PI positive cells was present in the *ΔmlaE ΔpqiA ΔyebS* mutant cultures, suggesting that the IM remains intact (Figure S1D). Overall, these results are consistent with a more severe OM defect in the triple mutant, suggesting that the Mla, Pqi, and Yeb systems act in concert to maintain the integrity of the OM barrier.

To better understand the nature of the OM defect in these mutants, we examined intact *ΔmlaE ΔpqiA ΔyebS* triple mutant *E. coli* cells by cryo transmission electron microscopy. Strikingly, most triple mutant *ΔmlaE ΔpqiA ΔyebS* cells exhibited cell surface ruffling, suggestive of gross disorganization of the OM and periplasm (Figure 1D), whereas the IM and OM of wild-type cells were smooth with a well-defined periplasm in between (Figure 1D). This is consistent with results from *S. coelicolor* in which mutations in the single MCE gene cluster led to global changes in the cell surface (Clark et al., 2013). Thus, MCE proteins are important for maintaining the morphology of the OM and the uniform spacing between the IM and OM.

### MCE protein MlaD forms a hexameric ring with a central hydrophobic pore

As MCE proteins have no detectable similarity to proteins of known structure or function, we obtained structural information using a combination of X-ray crystallography and electron microscopy (EM). MlaD is predicted to be anchored to the IM by a single N-terminal transmembrane helix, with its MCE domain residing in the periplasm (Figure 2A). We determined the crystal structure of the complete periplasmic domain of *E. coli* MlaD (residues 32-183) at 2.85 Å resolution and its core MCE domain at 2.15 Å resolution (Tables S1 and S2; Figure S2B). The MCE core domain adopts a 7-stranded β-barrel fold, encompassing just over 100 residues (Figures 2B and 2C). Most of the connecting loops are fairly short, with the exception of the β6-β7 loop, which projects prominently from one end of the barrel (Figure 2B). A search of the Protein Databank for structural homologs using DALI (Holm and Rosenström, 2010) did not identify any other proteins that share this fold, although we identified two classes of similar domains that differ in the arrangement of one or more β-strands (Figure S2A).

**Figure 2.**
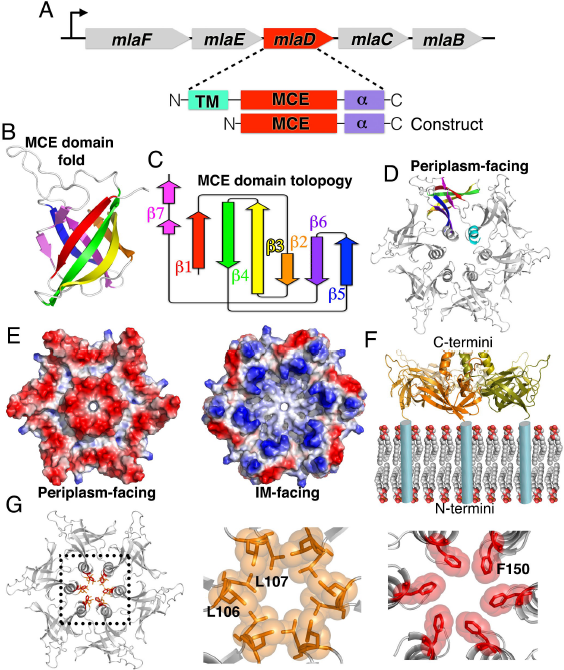
MCE protein MlaD forms hexameric rings with a central hydrophobic pore. (A) Schematic of *E. coli mla* operon and architecture of full-length MlaD and “periplasmic domain” construct. (B) Crystal structure of core MCE domain of MlaD reveals a novel, 7-stranded beta-barrel fold.
(C) Topology diagram of core MCE domain of MlaD [adapted from diagram generated by Pro-Origami (Stivala et al., 2011)]. (D) 6 copies of MlaD come together to form a homo-hexameric ring. (E) left, Electrostatic potential surface of MlaD in the same orientation as in (D), illustrating the net negative charge on the C-terminal/periplasm facing side. Right, in contrast, the membrane facing, N-terminal face of the MlaD ring is more positively charged. (F) 90˚ rotation of MlaD from depiction in (D) reveals that the N-terminus of each MCE domain projects from one face of the MlaD ring, docking this face of the ring in close proximity to the periplasmic face of the IM via the N-terminal TM helix from each subunit. (G) Two rings of hydrophobic resides gate the central channel of the MlaD ring. Middle, Leu106 and Leu107 at the tip of the β5-β6 loop. Right, Phe150 from the C-terminal helical bundle.

The six copies of MlaD are assembled into a homo-hexameric ring with the C-terminal α-helical domain extending from one face of the ring (Figure 2D). The interface between subunits consists of two main parts. First, the core MCE domains interact in a “head-to-tail” fashion, mediated primarily by the connecting loops at each end of the β-barrel fold, accounting for more than half (∼900 Å2) of the total interface (1,660 Å2). Second, the C-terminal helical regions from each protomer come together to form a 6-helix bundle at the center of the ring (760 Å2 interface). Truncation of the C-terminal α-helical region led to the formation of heptameric instead of hexameric rings (Figure S2C–S2G) suggesting that interactions between the C-terminal α-helices may play a role in determining the overall oligomeric state and packing of the MCE subunits.

The N-terminus of each MCE domain is connected by a short peptide linker to an N-terminal transmembrane helix, which places the N-terminal face of the ring in close proximity to the negatively charged phospholipid head groups (Figure 2F). Consistent with this orientation, the N-terminal, membrane-facing surface of the ring is considerably more basic than the C-terminal, periplasm-facing side (Figure 2E). The C-terminal helix from each subunit associates to form a hollow barrel very similar in structure to computationally designed 6-helix coiled-coils (Thomson et al., 2014; Zaccai et al., 2011), leaving a solvent accessible channel running through the center of the ring (Figure 2G). This channel is lined with hydrophobic residues and has two notable constriction points: the first is formed by Leu106 and Leu107 at the tip of the β5-6 loop and second by Phe150 in the C-terminal helix (Figure 2G). These narrowings, in particular the constriction formed by the Phe150 ring, are reminiscent of the Phe427 Φ-clamp in the central pore of anthrax protective antigen (PA), where they play a key role in the transport of protein toxins through the PA translocon (Jiang et al., 2015). At the narrowest point, created by Leu106 and Leu107 at the tip of the β5-6 loop, the channel diameter is ∼6 Å, large enough to allow the passage of small hydrophobic molecules like phospholipids or sterols. It should also be noted that the weaker electron density in this region of the crystal structure suggests that these loops are at least somewhat flexible and therefore may allow somewhat larger molecules to pass through. Based upon its orientation relative to the IM (Figure 2F), the MlaD ring is poised to function as a channel to facilitate trafficking of small hydrophobic molecules to or from the inner membrane.

### Architecture of the MlaFEDB transporter complex

An MCE protein implicated in lipid transport in plants, TRIGALACTOSYLDIACYLGLYCEROL 2 (TGD2), forms part of a high-molecular weight complex in the chloroplast inner membrane (Awai et al., 2006; Roston et al., 2011, 2012). To assess whether MlaD forms a complex with any of the other Mla proteins, we over-expressed the full *mlaFEDCB* operon in *E. coli* (Figure 3A), solubilized the membrane fraction with detergent, and determined whether any other Mla proteins co-purify with His-tagged MlaD. After affinity purification and gel filtration (Figure 3B), SDS-PAGE analysis of the purified MlaD sample revealed the presence of three additional proteins forming a stable complex with MlaD (Figure 3C). Excision of these bands and protein identification by mass spectrometry revealed that these proteins were MlaF (an ABC-type ATPase), MlaE (a multi-pass transmembrane protein and likely permease), and MlaB (STAS domain containing protein); the co-expressed MlaC protein, however, was absent. Independently, similar results were recently reported (Thong et al., 2016).

**Figure 3.**
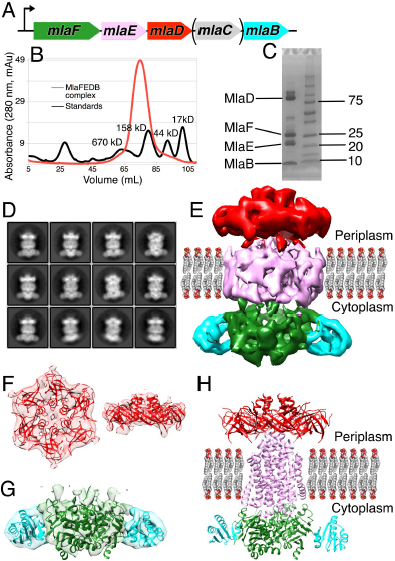
MlaD is part of a large transport complex in the IM. (A) Co-expression of the *mlaFEDCB* operon with 6xHis tagged MlaD. (B) Gel filtration of detergent solubilized MlaD yields a high molecular weight complex. (C) SDS-PAGE analysis of complex and mass spectrometry reveals a multi-protein complex including MlaF, MlaE, MlaD, and MlaB. Sample was not boiled to prevent aggregation of membrane proteins, so apparent molecular weight of protein bands are unreliable. Only MlaC is not present. (D) 2D cryo-EM class averages for the MlaFEDB complex are feature rich, including possible transmembrane helices and clearly discernible densities for the MlaD ring and V-shaped, dimeric ABC ATPase subunits. (E) 3D reconstructed volume of MlaFEDB complex in the IM at ∼10 Å resolution, with large periplasmic and cytoplasmic domains. Density segmented and colored to indicate locations of MlaD (red), MlaE (pink), MlaF (green), and MlaB (cyan). (F) Left, top view (looking down at complex from the OM towards the IM) and right, side view (parallel to the plane of the IM) of 6-fold symmetric periplasmic domain with hexameric MlaD crystal structure fit into the map. (G) Side view (parallel to the plane of the membrane) of the 2-fold symmetric cytoplasmic domain, with homology models of the MlaF ATPase (green) and MlaB STAS domain (cyan) subunits fit into the map. Note that the orientation of MlaB relative to MlaF is not well defined by the map. (H) hybrid model of the MlaFEDB complex, including MlaD crystal structure (MCE subunit, red), homology models of MlaF (ATPase, green) and MlaB (STAS, cyan) subunits, and a *de novo*, computationally predicted model for the MlaE permease subunit (pink)(Ovchinnikov et al., 2015).

We next sought to determine the architecture of the purified MlaFEDB complex by single particle cryo-electron microscopy (cryo-EM). The 2D class averages were feature-rich, and individual components within the complex were clearly identifiable (Figure 3D). Some inherent flexibility in the complex was apparent in these 2D class averages, suggesting that the relative orientation of the components is not rigidly fixed, which may also limit the final resolution of our density maps. 3D classification and refinement produced a ∼10 Å resolution density map (Figures 3E, S3A, and S3B). The six-fold symmetry of the MlaD ring is readily apparent at one end of the map (Figure 3F), which could be well fit by the hexameric, MlaD crystal structure (Figure 3F), with additional density for the N-terminal transmembrane helix from each subunit extending down towards the membrane region. The opposite end of the map is approximately two-fold symmetric, consistent with the dimeric state of known ABC transporters. Homology models of MlaF and MlaB, based upon clear similarity to ABC ATPases and STAS domain proteins, respectively, were generated (Kelley et al., 2015) and were used to fit into the density. Two copies of the MlaF ATPase model can be fit into the map (Figure 3G), leaving two small densities on adjacent to each MlaF subunit. These two densities are approximately the size and shape of a STAS domain, allowing for the docking of two copies of the MlaB model into the map (Figure 3G). It should be noted that while the overall position of MlaB within the complex is clear, its orientation relative to MlaF is not well defined by the map and the details of this interaction await further investigation. The remaining central density most likely corresponds to the transmembrane subunit, MlaE, along with six additional transmembrane helices from the MlaD subunits. Based upon the volume of the transmembrane region and the apparent 2-fold symmetry of the map, we infer that there are most likely two copies of the MlaE subunit, although the modest resolution of our cryo-EM reconstruction does not allow us to trace the MlaE subunit *de novo*. The apparent symmetry mismatch between the hexameric MlaD and the dimeric permease assemblies implies that the MlaD-MlaE interface cannot be identical for each of the six MlaD subunits.

Recently, a model for the MlaE structure was generated from a powerful new approach to protein structure prediction using residue contact restraints based upon co-evolving residue pairs (Ovchinnikov et al., 2015). This *de novo* model for MlaE is dimeric and fits within the density of the transmembrane region (Figure 3E and 3H). Additional density surrounds the MlaE model, which may be derived from the six MlaD transmembrane segments associating with the sides of MlaE within the transmembrane region, as well as the amphipol used to solubilize the complex. Indeed, several residues in the MlaD transmembrane helices show signatures of co-evolution with residues in the membrane spanning portion of MlaE, suggesting a physical interaction between these regions (Figure S3C). Several additional residues in the MlaF ATPase appear to be co-evolving with residues in MlaE. These residues lie at the end of MlaE predicted to be oriented towards the cytoplasm, and on the surface of MlaF typically used by other ABC ATPases to interact with the transporter permease subunit (Figure S3D), supporting the placement and orientation of the Mla subunits in our cryo-EM map. Strikingly, this predicted model of the MlaE permease subunit closely resembles the overall transmembrane fold of ABCG5/ABCG8 (Figure S3E), a recently determined human ABC transporter structure involved in the export of cholesterol from hepatocytes (Lee et al., 2016). While the structure of MlaE remains to be determined experimentally, true homology with ABCG5/ABCG8 would group MCE transporters with this important class of ABCG transporters exporting another hydrophobic substrate, cholesterol, in humans.

### Structure of MlaC shows that it binds lipid tails

Many ABC transporters from double-membraned bacteria employ small substrate binding proteins that facilitate substrate recognition and trafficking across the periplasm. Although MlaC has no sequence similarity to the conventional families of substrate binding proteins, it is part of the *mla* operon (Figure 4A), and targeted to the periplasm by an N-terminal signal sequence. Therefore MlaC may play a role in shuttling substrates to or from the MlaFEDB transporter complex. The structures of two possible homologs of MlaC were previously deposited by the Northeast Structural Genomics Consortium (NESG) without any associated publications (2QGU and 4FCZ; 27% and 22% sequence identity and Cα rmsd of 2.5 Å and 3.0 Å with *E. coli* MlaC, respectively). The NESG crystal structure of an MlaC-like protein from *R. solanacearum* (2QGU) was also crystallized with a ligand of unknown origin bound in its hydrophobic pocket, which was modeled as phosphatidylethanolamine (Figures S4D and S4G). Based upon these structures, it has been proposed that MlaC-like proteins may function as periplasmic substrate binding proteins for the transport of lipids (Malinverni and Silhavy, 2009) or other hydrophobic molecules.

**Figure 4.**
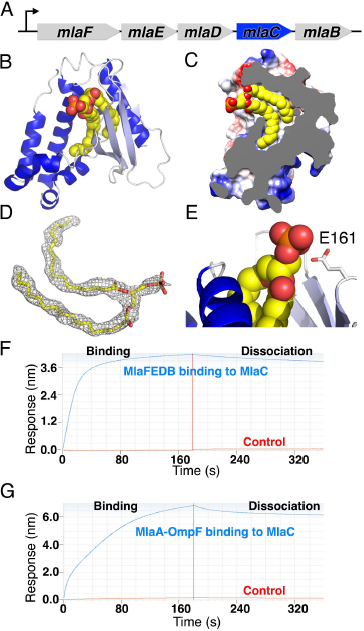
MlaC is a periplasmic protein that shuttles lipids across the periplasm. (A) MlaC is encoded in the *mla* operon and is the only component not stably present in the MlaFEDB complex. (B) Crystal structure of MlaC adopts a mixed α/β fold with 4 β-strands (light blue) and 7 α-helices (dark blue). MlaC-bound lipid shown as spheres. (C) Cut away view of MlaC surface, revealing deep hydrophobic pocket in MlaC containing the two acyl chain “tails” of a bound lipid. (D) Unbiased 2Fo-Fc electron density map for bound lipid (prior to refinement with lipid ligand), contoured at 1 sigma. (E) In contrast to the extensive interactions between MlaC and the lipid tails, no significant interactions are made between MlaC and the lipid head group. (F and G) MlaC interacts with the IM, MlaFEDB complex (F) and MlaC also interacts with the OM MlaA-OmpF complex (G) by biolayer interferometry, suggesting MlaC may ferry lipid substrates between the IM and OM components of the Mla transport system.

To better understand how MlaC might be involved in substrate recognition, we determined the crystal structure of MlaC at ∼1.50 Å resolution. Like 2QGU and 4FCZ, MlaC adopts a mixed α/β-fold, consisting of a highly twisted, 4-stranded β-sheet and a bundle of 7 α-helices (Figure 4B). Outside of this small group of three MlaC-like proteins, a structure-based comparison to other proteins in the PDB using DALI revealed more distant similarities to the nuclear transport factor 2 (NTF2) protein superfamily and to the parental, Cystatin-like fold (Figure S4A). Interestingly, within a deep hydrophobic pocket in the protein core of MlaC, we observed a roughly “horse-shoe”-shaped density (Figure 4D) that closely resembles two fatty acid chains esterified to a phosphoglycerol head group, which we have modeled as phosphatidic acid (Figures 4B, 4C, S4C, and S4F). The two acyl chains of the lipid insert snugly into the protein core where they are completely protected from solvent in the protein core (Figure 4C), while the head group is almost entirely solvent exposed and makes only minor contacts with the protein (Figure 4E). Due to the limited volume of the hydrophobic cavity and the lack of head group interactions, we predict that MlaC is capable of binding to a wide range of diacyl lipids with little head group specificity. In another MlaC-like structure in the PDB (4FCZ), no ligands were explicitly modeled, but our re-refinement of the deposited coordinates and structure factors revealed additional electron density in the hydrophobic pocket of this protein (Figures S4E and S4H, and Table S3), suggestive of a tetra-acyl, cardiolipin-like lipid. The observation of lipid ligands in MlaC, 2QGU, and 4FCZ support the notion that MlaC and its close homologs are lipid-binding proteins.

### MlaC binds IM and OM protein complexes

Since MlaC is a soluble lipid-binding protein predicted to localize to the periplasm, we hypothesized that MlaC may shuttle lipid substrates across the periplasm between the IM MlaFEDB transporter and an OM complex, where lipids would be inserted into or removed from the OM. Lipoprotein MlaA (which is not part of the *mla* operon in *E. coli*) has emerged as candidate component for this OM complex, as it has been implicated in Mla function (Malinverni and Silhavy, 2009) and forms a complex with the OM porin proteins, OmpC and OmpF (Chong et al., 2015). Thus, we set out to determine whether MlaC could interact directly with purified MlaFEDB and MlaA-OmpF protein using biolayer interferometry. We over-expressed and purified MlaA from *E. coli*, which co-purified as a stable complex with OmpF under our culture conditions, as confirmed by mass spectrometry. We observed robust and specific binding of both MlaFEDB (Figure 4F) and MlaA-OmpF assemblies (Figure 4G) to MlaC, as well as a direct interaction between MlaC and MlaD (Figure S4I). These results demonstrate that MlaC is capable of binding to both the IM and OM complexes, supporting a model where MlaC plays a central role in transferring a lipid substrate from one membrane to another.

### Stacking of MCE rings yields larger assemblies in YebT and PqiB

No other homologs of MlaC are encoded in the *E. coli* genome, raising the question of how the Yeb and Pqi systems might move substrates across the periplasm. To gain insight into this question, we purified full-length *E. coli* YebT and the periplasmic domain of PqiB, which contain seven and three tandem MCE domains, respectively (Figure 5A and 5D), and obtained negative-stain EM reconstructions at ∼20 Å resolution (Figures 5B, 5C, 5E, 5F, S5A, and S5B).

**Figure 5.**
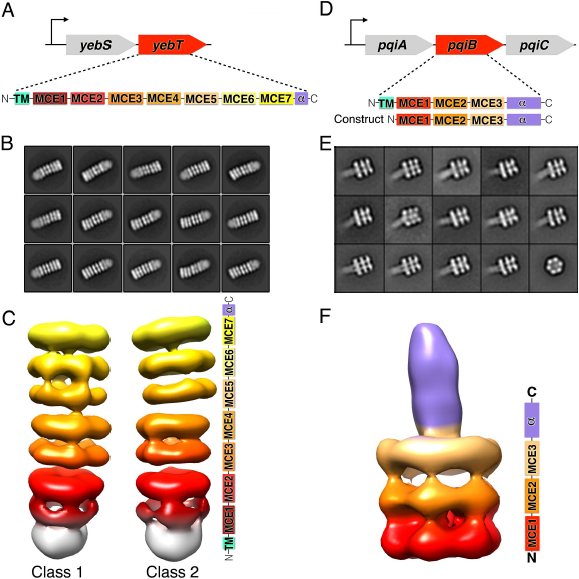
YebT and PqiB adopt elongated tube and syringe-like architectures. (A) *E. coli yeb* operon and domain architecture of MCE protein YebT. (B) Negative stain, 2D class averages for YebT. (C) 3D reconstructions of YebT at ∼20 Å resolution, revealing a 7-layered barrel formed from a stack of 7 MCE rings (colored shades of red, orange, and yellow). At right, domain organization of YebT as it relates to the EM electron density map. The map and schematic are color coded to indicate how the seven tandem MCE domains (colored shades of red, orange, and yellow) contribute to the seven stacked hexameric rings in the final, homohexameric assembly. A small additional density at one end (white), corresponds to the transmembrane regions in the IM. Reconstructions are viewed from the side, parallel to the plane of the IM, with the IM towards the bottom and the OM towards the top. (D) *E. coli pqi* operon and domain architecture of MCE protein PqiB, including “periplasmic construct” used for structural studies. (E) Negative stain, 2D class averages for PqiB, revealing a 3-layered barrel with a “needle” protruding from one end. (F) 3D reconstruction of PqiB at ∼20 Å resolution, viewed from the side (similar to YebT), with domain organization at right.

The overall architecture of YebT is striking, consisting of a tall stack of 7 rings of density forming an elongated, tube-like barrel, ∼230 Å long and ∼90 Å in diameter (Figure 5C). To a first approximation, YebT resembles other macromolecular barrels such as the proteasome, ClpXP, and GroEL, wherein one or more small globular domains assemble into rings that in turn stack on top of one another (Figure S5F). However, the YebT barrel is much longer than any of these other examples, both in terms of number of stacked rings (7 for YebT vs 4 each for the proteasome and GroEL), and absolute length (∼230 Å for YebT vs 150 Å and 135 Å for the proteasome and GroEL, respectively). An additional, fuzzy density is observed at one end of the barrel, which likely corresponds to a detergent micelle and the N-terminal transmembrane helices from each YebT polypeptide chain (Figure 5C, bottom).

In contrast to the 7-ringed tube of YebT, the negative stain EM reconstruction of PqiB resembles a needle and syringe, with the syringe barrel consisting of a stack of three rings of density (Figure 5F). A needle-like projection emanates from the center of the top of the barrel. Recombinant PqiB proteins containing only the three MCE domains form barrels without needles, showing that the C-terminal helical regions from each subunit come together to form this needle-like structure (Figures S5C and S5D).

Each of the rings of density in the YebT and PqiB reconstructions exhibit apparent 6-fold symmetry, and is similar in overall size and shape to the hexameric ring formed by the MCE domains of MlaD. Indeed, polyalanine MCE models derived from the MlaD hexamer fit well into each of the density rings (Figure S5E). Thus, we propose that YebT is a ∼570 kDa hexameric assembly of 6 copies of the YebT polypeptide which is anchored in the IM via 6 transmembrane helices in a manner analogous to MlaD. In this orientation, IM distal end of YebT could be as much as ∼230 Å away from the inner membrane. Similarly, hexameric PqiB would form a ∼360 kDa complex, anchored on the inner membrane by the face of one ring, with the syringe barrel and needle projecting up to ∼230 Å away from the IM. As the distance between the IM and OM in *E. coli* is approximately 225 Å (Hoffmann et al., 2008; Matias et al., 2003), this raises the possibility that YebT and PqiB might directly span the intermembrane space, facilitating the transport of substrates (likely lipids or other hydrophobic molecules) across the intervening, aqueous periplasm.

### Cryo EM structure of PqiB

In order to better understand how these larger MCE domain proteins such as YebT and PqiB may be involved in lipid transport across the periplasm, we used single-particle cryo EM to obtain a reconstruction of the PqiB periplasmic domain with an overall resolution of 3.96 Å (Figures S6A - S6C). However, much of the structure was resolved at significantly higher resolution, with the local resolution across much of the MCE domain barrel in the vicinity of ∼3.1 Å (Figure 6A). The final maps are of high quality, with nearly continuous backbone density across the three MCE rings and rich side-chain density which allowed the assignment of the amino acid sequence over majority of the final model (Figure S6D).

**Figure 6.**
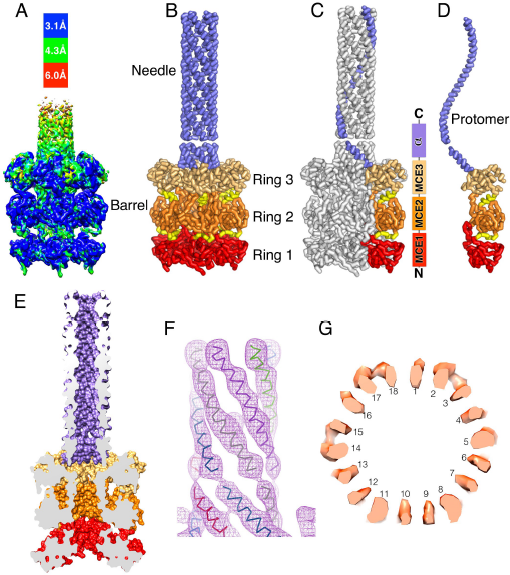
Cryo EM structure of PqiB reveals architecture of a periplasmic bridge. (A) Cryo EM map for PqiB with an overall resolution of 3.96 Å. Map is colored by resolution, indicating that most of the MCE rings forming the barrel have a local resolution close to ∼3.1 Å. (B) Model of PqiB periplasmic domain derived from cryo EM reconstructions. PqiB is viewed from the side, parallel to the plane of the IM, with the IM towards the bottom and the OM towards the top. The MCE1, MCE2, and MCE3 rings of the barrel are colored red, orange, and tan, respectively, with inter-ring connectors colored yellow. The needle, derived from the C-terminal helical regions, is colored lilac. (C and D) Similar views as in (B), but with only one subunit colored in the complex (C), and in isolation (D). (E) Surface representation of PqiB model, cut in half vertically to reveal a long, continuous tunnel running the full length of the needle and barrel. (F) Helical density for the first half of the needle, proximal to the barrel. The distal half was modeled as a continuation of this 6-helix coiled-coil structure. (G) Horizontal slice through the PqiB barrel between rings 2 and 3, revealing 18 tubes of density resembling 18 β-strands. Similar density is observed between rings 1 and 2, suggesting that the central pore through the MCE rings is lined by an 18-stranded, β-barrel like structure.

As suggested by our lower resolution negative-stain reconstructions, six PqiB polypeptides associate to form the complete, ∼360 kDa PqiB needle-and-syringe assembly (Figure 6B). The needle of PqiB is formed by a hollow, 6-helix coiled-coil, while the barrel consists of a total of 18 MCE domains, distributed into 3 stacked hexameric rings (Figures 6B - 6D). The MCE1, MCE2, and MCE3 domains from the primary sequence each self-associate with the same domain from other PqiB polypeptide chains to form each of the three observed rings: the MCE1 ring at one end of the barrel; the MCE2 ring in the middle of the barrel; and the MCE3 ring on the other side of the barrel, immediately adjacent to the base of the needle (Figure 6C). Consequently, the number of stacked rings in a given MCE protein seems likely to be determined by the number of tandem MCE domains in the protein. This is also the case for YebT, in which seven MCE domains in the primary sequence yield an architecture consisting of seven rings (Figure 5C).

PqiB is hollow in the center and open at either end, consistent with a possible function as a pore for the translocation of lipids (Figure 6E). In contrast to some other macromolecular barrels, like the proteasome, where a staggered arrangement of the domains in the stacked rings helps to fill in the gaps between subunits in the barrel wall, the three MCE domains from each polypeptide chain sit roughly one on top of the next in a straight line (Figure 6D). Because of this linear arrangement, the walls of the barrel formed by the core the MCE domains themselves are incomplete, with large gaps 10-20 Å in diameter between the rings. However, in these two regions - between the MCE1 and MCE2 rings, and between the MCE2 and MCE3 rings - the central pore of PqiB appears to be lined by an 18-stranded β-barrel motif (Figure 6G), creating a barrier that could exclude solvent from a hydrophobic, lipid conducting channel (see Method Details in Supporting Information).

At the C-terminus of PqiB's final MCE domain, there is clear additional density for 6 helices (Figure 6F), one contributed from each polypeptide chain, forming an extended 6 helix coiled-coil. The resulting needle has an outer diameter of ∼35 Å and a hollow lumen approximately 15-20 Å wide that is continuous with the channel running through the PqiB (Figure 6E). A similar arrangement of 6 helices was observed near the C-terminus of MlaD in our crystal structure, though the twist of the coiled-coil is more pronounced in PqiB, resulting in a wider pore through the PqiB needle. After ∼ 20 helical residues, there is a pronounced break in the density followed by an additional ∼40 helical residues, in good agreement with the predicted secondary structure for this region and a helix-breaking proline residue near the junction between these two helical segments (Figure 6F). The needle is somewhat flexible, as is clear from the 2D class averages (Figure S6B), leading to weaker density for the needle as the distance increases from the barrel, and beyond the first ∼60 helical residues, the density becomes uninterpretable. However, most of the remaining 50-60 residues in C-terminal region of PqiB are predicted to be helical, and it seems likely that the observed 6-helix coiled-coil structure continues to nearly the C-terminus. This would result in a needle approximately 135 Å long, in good agreement to the maximum needle lengths we observed in unaveraged, single particle images. From the bottom of the barrel to the tip of the needle, this would make PqiB ∼230 Å long, similar to the length of YebT and the ∼225 Å distance between the IM and OM. Thus, both PqiB and YebT are poised to serve as direct conduits for the transport of lipids or other hydrophobic molecules directly across the periplasm, without the need for a soluble carrier protein like MlaC.

## DISCUSSION

### Model for lipid transport by MCE systems

These structural data allow us to generate a model for how MCE transport systems traffic lipids across the periplasm (Figure 7). We present this model in the context of lipid import to the IM. However, as the direction of transport has not been absolutely established, these systems could also drive lipid import towards the inner membrane, with the steps outlined occurring in the reverse order. In the case of the Mla system, lipids for import are extracted from the outer membrane by the MlaA-OmpC/OmpF complex, perhaps via a lateral pathway into the membrane through the opening of the β-barrel structure of OmpC/OmpF, as has been proposed for the LptD subunit of the LPS transporter. Binding of MlaC to the lipid-bound MlaA-OmpC/OmpF complex results in the transfer of the lipid to the hydrophobic pocket of MlaC, where it is protected from solvent. Lipid bound MlaC can then diffuse across the periplasm to the MlaFEDB complex in the IM, where ATP hydrolysis may facilitate extraction of the lipid from MlaC and/or the translocation of the lipid into the IM. In contrast, the segmented tubes formed by YebT and the syringe-like architecture of PqiB are poised to bridge the gap between the IM and OM directly, forming a continuous lipid translocation pathway that transverses the entire periplasmic space without the need for a soluble lipid carrier protein analogous to MlaC (Figure 7). This would be more similar to the model proposed for the transport of the other major lipid component of the OM, LPS, which is trafficked across a periplasm-spanning bridge comprised of the LptA, LptC, and LptD proteins (Figure 7) (Okuda et al., 2012; Ruiz et al., 2008; Sperandeo et al., 2007, 2008; Suits et al., 2008; Tefsen et al., 2005; Wu et al., 2006). However, in contrast to the open groove utilized by the LPS transport system, we propose that MCE transporters translocate lipids through a closed channel formed by the pores of multiple stacked MCE rings (e.g., YebT), or a combination of rings and a hollow, needle-like projection formed by long C-terminal helices (e.g., PqiB, and the MCE proteins from *M. tuberculosis* and chloroplasts; Figure S7B and S7C). Thus, at least three unique strategies appear to have evolved for lipid transport between the IM and OM of double-membraned bacteria and chloroplasts. First, a soluble lipid carrier protein such as MlaC can be employed, with a hydrophobic pocket accepting the lipid tails, protecting them from bulk solvent. This parallels the mechanism used by the Lol pathway to transport nascent lipoproteins to the outer membrane (Konovalova and Silhavy, 2015), in which the structurally unrelated protein LolA plays a similar functional role to MlaC by sequestering lipid modifications during the transport of nascent lipoproteins across the periplasm. Second, lipids can be translocated along a hydrophobic groove on a bridge between the IM and OM, as in the LPS transporter. And third, lipids may be transported through the interior of elongated, closed tunnels formed by large MCE domain proteins, as in PqiB and YebT. While MCE transporters appear to have evolved independently, this final strategy is mechanistically similar to the approach employed by bacterial efflux pumps such as the AcrA/AcrB/TolC system, which create a periplasm-spanning tunnel to transport hydrophobic antibiotics, dyes, and other toxic molecules across the periplasm to the exterior of the cell (Du et al., 2014).

**Figure 7.**
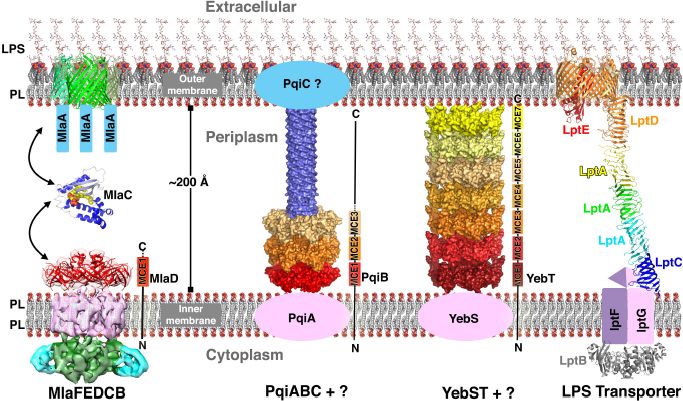
Model for MCE-mediated phospholipid transport across the periplasm in *E. coli*. Left, model for the Mla lipid transport system. MlaC likely serves as a soluble lipid carrier protein to transport lipids between the OM MlaA-OmpC/OmpF complex and the IM MlaFEDB complex. Middle left, the MCE protein PqiB is significantly larger than MlaD, with three stacked rings of MCE domains and a long projecting helical domain. A predicted OM lipoprotein, PqiC, may interact with the tip of the PqiB needle while PqiA may associate with the MCE1 ring at the inner membrane. These additional IM and OM factors may mediate lipid extraction from or insertion from the membranes. Middle right, YebT has seven stacked MCE domain rings, and may associate with YebS in the IM. Both PqiB and YebT are sufficiently long to bridge the aqueous periplasm and may mediate the transport of lipids directly between the IM and OM through the central pores of the stacked MCE rings and needle-like 6-helix coiled-coils (without a soluble carrier resembling MlaC). Right, model of LPS transport system, which has also been suggested to form a bridge across the periplasm, but uses a structurally unrelated framework to create a continuous lipid transport path.

Many important questions relating to the function of MCE transporters remain to be explored. In the case of LPS transport, the IM LptBFG complex uses ATP hydrolysis to extract LPS molecules from the IM and transport them through the LptACD bridge to the OM. It is conceivable that the Mla, Pqi, and Yeb systems may function in a similar way, using ATP hydrolysis to drive phospholipid export to create new membrane as the cell grows, perhaps in conjunction with other transport mechanisms. Alternatively, MCE systems may be involved in retrograde transport of mislocalized phospholipids from the OM to the IM (Chong et al., 2015; Malinverni and Silhavy, 2009; Sutterlin et al., 2016; Thong et al., 2016), or for the import of of other hydrophobic molecules to be used for energy (Klepp et al., 2012; Mohn et al., 2008; Pandey and Sassetti, 2008) or as precursors in biosynthetic processes (Awai et al., 2006; Roston et al., 2011, 2012, Xu et al., 2003, 2008). Based on our understanding of the Mla system, it is likely that the periplasm spanning components YebT and PqiB will link proteins in the IM and OM to facilitate transport of their substrates. A small lipoprotein encoded immediately downstream from *pqiB*, called PqiC (aka, YmbA), maybe be part of an outer membrane complex interacting with PqiB, though no homologous gene is present in the *yeb* operon. Similarly, the predicted integral membrane proteins YebS and PqiA, encoded just upstream of *yebT* and *pqiB*, and apparently structurally different from MlaE, may form part of IM complexes for each system, though ATPase genes are lacking from both the *yeb* and *pqi* operons. While the corresponding ATPase subunits may be be encoded elsewhere in the genome, genetic evidence suggests that the MlaF ATPase is not shared between all three systems (data not shown), and we also cannot rule out the possibility that the Yeb and Pqi systems may be driven by proton motive force, or even facilitate passive equilibration of lipids between the IM and OM rather than active transport. Further studies will be required to identify additional components that may function together with YebST and PqiABC.

The hexameric organization of the *E. coli* MCE proteins provides a possible explanation for a puzzling observation in mycobacterial species such as *M. tuberculosis*, where six distinct MCE genes are usually grouped together in an operon (Figure S7A) (Casali and Riley, 2007): it seems likely that MCE proteins in these species will form hetero-hexamers. However, it remains unclear why *M. tuberculosis*, *E. coli*, and many other organisms encode multiple MCE systems. While they may serve partially redundant roles, their very distinct structures suggest they may have distinct substrate specificities, transport lipids in different directions, or might be subject to unique regulatory control. Intriguingly, despite the fact that MCE proteins are almost universally conserved from double-membraned bacteria to chloroplasts, we have been unable to identify homologs in mitochondria, and it is unclear if this reflects extreme drift of the protein sequences in the ∼1 billion years since the divergence of bacteria and mitochondria, or if they have been functionally replaced during domestication of this organelle. Future studies will begin to unravel these and other questions to further understand the assembly and maintenance of the outer membranes of bacteria and organelles.

## AUTHOR CONTRIBUTIONS

D.C.E. and G.B. conceived the research; D.C.E., G.B., and G.G. collected data; D.C.E., G.B., R.V., and S.O. analyzed data; D.C.E., G.B., and R.V. wrote the manuscript. D.C.E., G.B., G.G., S.O., J.S.C, and R.V. edited the manuscript.

## ACKNOWLEDGEMENTS

We thank C. Waddling from the UCSF Crystallography and SAXS Facility for support with crystallization; J. Holton, G. Meigs, and the staff of Advanced Light Source beamline 8.3.1 for beamline support; A. Gray, C. Gross, and M. Elvekrog (UCSF) for providing plasmids and bacterial strains; N. Stuurman and A. Williamson (UCSF) for assistance with fluorescence microscopy; G. Isom and I. Henderson (U. Birmingham), Y. Cheng, E. Palovcak, JP. Armache and other members of the Cheng lab (UCSF), and A. Brilot (UCSF) for helpful suggestions and discussions; M. Braunfeld for EM technical assistance; C. Kennedy for assistance with high performance computing; C. Hong and Z. Yu at Janelia Research Campus for EM data collection; the UC Davis Campus Mass Spectrometry Facilities for protein MS assistance; and M. Morrissey, X. Su, M. Tanenbaum, Y. Cheng, and I. Wilson for critical reading of the manuscript. This work was supported by NIH grants K99GM112982 (G.B.) and 1S10OD020054, and the Howard Hughes Medical Institute (R.D.V.). D.C.E. is a Damon Runyon Fellow supported by the Damon Runyon Cancer Research Foundation (Grant DRG-2140-12). EM data were collected at the HHMI CryoEM Shared Facility at Janelia Research Campus. The Advanced Light Source is supported by the Director, Office of Science, Office of Basic Energy Sciences, of the US Department of Energy under Contract DE-AC02-05CH11231.

## SUPPLEMENTAL INFORMATION

Supplemental Information includes seven figures, three tables, and can be found with this article at http://dx.doi.org/xxxxxxxxxx

## METHOD DETAILS

### Bacterial strains and growth conditions

All *E. coli* strains used in this work are derivatives of the wild-type strain MG1655. Mutant strains were constructed by P1 transduction of Kanamycin resistance-marked alleles from an appropriate strain from the Keio collection (Baba et al., 2006).

### TEM of *E. coli* outer membrane

Overnight cultures of MG1655 and mutant strains were diluted 1:100 in LB and grown with shaking at 37°C until an OD600 of ∼0.7. 3µl of culture was spotted onto an electron microscopy grid (Quantifoil 2/2), and cells allowed to attach for ∼30 seconds. The grid was blotted for 20 seconds and then plunge frozen in liquid ethane using an FEI Vitrobot Mark III. The grids were imaged using a FEI Polara microscope operating at 300keV, and equipped with a Gatan energy filter and K2 camera. UCSF tomography software was used to collect images of the wild-type and mutant cells.

### Expression and purification of MlaD

DNA corresponding to the “periplasmic domain” (residues 32-183) and “core MCE domain” (residues 32-140) of MlaD was amplified from the MG1655 genome and cloned by Gibson assembly (Gibson et al., 2009) into a custom pET vector resulting in an N-terminal 6xHis tag followed by a TEV protease cleavage site just upstream of MlaD. The resulting plasmids (pBE1160 and pBE1161, respectively) were transformed into Rosetta 2 cells (Novagen). For expression, overnight cultures of Rosetta 2 / pBE1160 or Rosetta 2 / pBE1161 were grown at 37˚ C with shaking to an OD600 of ∼0.9, then induced by addition of IPTG to a final concentration of 1 mM and continued incubation overnight shaking at 15˚ C. Cultures were harvested by centrifugation, and pellets were resuspended in lysis buffer (50 mM Tris pH 8.0, 300 mM NaCl, 10 mM imidazole). Cells were lysed by two passes through an Emulsiflex-C3 cell disruptor (Avestin), then centrifuged at 38,000g to pellet cell debris. The clarified lysates were incubated with NiNTA resin (Qiagen) at 4˚ C, which was subsequently washed with Ni Wash Buffer (50 mM Tris pH 8.0, 300 mM NaCl, 10 mM imidazole) and bound proteins eluted with Ni Elution Buffer (50 mM Tris pH 8.0, 300 mM NaCl, 250 mM imidazole). MlaD containing fractions eluted from the NiNTA column were pooled and concentrated before separation on a Superdex 200 gel filtration column (GE Healthcare) equilibrated in 20 mM Tris pH 8.0 and 150 mM NaCl.

### Crystallization and structure determination of MlaD

Gel filtration fractions containing purified MlaD proteins were concentrated to 20-60 mg/mL and sitting drop, vapor-diffusion crystallization trials were conducted using the JCSG Core I-IV screens (Qiagen). Crystals of the core MCE domain of MlaD (residues 32-140) grew from drops consisting of 100 nL protein plus 100 nL of a reservoir solution consisting of 0.1 M sodium acetate pH 4.5, and 40% 1,2 propanediol, and were cryoprotected by supplementing the reservoir solution with 25% ethylene glycol. Native diffraction data was collected at ALS beamline 8.3.1, and indexed to C2 and reduced using XDS (Table S1) (Kabsch, 2010).

As MlaD has no homology to proteins of known structure, and has no Met residues in the core MCE domain, we set out to generate a panel of single mutants in which each Leu residue of the core domain was converted to Met (Leu42Met, Leu52Met, Leu73Met, Leu79Met, Leu84Met, Leu99Met, Leu106Met, Leu107Met, Leu112Met, Leu114Met, Leu123Met, and Leu128Met). Of these 12 target mutants, we successfully generated 11 by site directed mutagenesis (reaction for Leu73Met failed). Test expressions revealed that 5 of these 11 mutants expressed at near wild-type levels (Leu79Met, Leu84Met, Leu106Met, Leu107Met, Leu123Met). We then prepared selenomethionine-derivatived protein for each of these five mutants and screened for diffraction-quality crystals, which were obtained for 4 of the 5 mutants. Multi-wavelength anomalous dispersion (MAD) datasets were collected from selenomethionine-derivatived crystals of the Leu84Met and Leu107Met mutants at ALS beamline 8.3.1. Anomalous signal from the Leu107Met crystals was weak despite high data quality and strong Se signal in X-ray fluorescence scans, and later this residue was found to be at the tip of a flexible/partially disordered loop. However, Leu84Met yielded a high quality data set, which was indexed to C2 and reduced using XDS (Kabsch, 2010), and phased by MAD using the SHELX C/D/E pipeline (Sheldrick, 2010) (Table S1 and S2). The polyalanine model from SHELXE was rebuild using the Autobuild module (Terwilliger et al., 2008) of Phenix. The resulting model was adjusted in Coot (Emsley et al., 2010) and refined using Phenix (Afonine et al., 2012). The final model consists of 7 copies of the MCE domain forming a hepatmeric ring.

Crystals of the complete MlaD periplasmic domain (residues 32-183) grew from drops consisting of 100 nL protein plus 100 nL of a reservoir solution consisting of 0.2 M zinc acetate, 0.1 M MES pH 6.0, and 15% ethanol, and were cryoprotected by supplementing the reservoir solution with 20% ethylene glycol. Native diffraction data was collected at ALS beamline 8.3.1, and indexed to C2 and reduced using XDS (Table S1) (Kabsch, 2010). This data set was phased by molecular replacement using Phaser (McCoy et al., 2007), with the MlaD core MCE domain as a search model. The resulting model was adjusted in Coot (Emsley et al., 2010) and refined using Phenix (Afonine et al., 2012). This final model consists of 3 copies of the MlaD periplasmic domain. Application of the 2-fold symmetry operator results in the biologically relevant hexameric ring.

### Purification of the MlaFEDB complex

DNA corresponding to the complete *mlaFEDCB* operon (from MlaF start codon to MlaB stop codon) was amplified from the MG1655 genome and cloned by Gibson assembly (Gibson et al., 2009) into a custom pET vector resulting in an C-terminal 6xHis tag. The resulting plasmid (pBE1200) was transformed into Rosetta 2 cells (Novagen). For expression, overnight cultures of Rosetta 2/pBE1200 were grown at 37˚ C with shaking to an OD600 of ∼0.9, then induced by addition of IPTG to a final concentration of 1 mM and continued incubation for ∼4hr shaking at 37˚ C. Cultures were harvested by centrifugation, and pellets were resuspended in lysis buffer (50 mM Tris pH 8.0, 300 mM NaCl, 10 mM imidazole). Cells were lysed by two passes through an Emulsiflex-C3 cell disruptor (Avestin), then centrifuged at 35,000g to pellet cell debris. The clarified lysates were transferred to ultracentrifuge tubes and centrifuged at 200,000g to pellet membrane fraction. Membrane pellets were resuspended in lysis buffer, then n-Dodecyl-β-D-Maltopyranoside (DDM; Anatrace) was added to a final concentration of 25 mM. After incubation for ∼1hr with rocking at 4˚ C, the DDM solublized membrane fraction was centrifuged at 200,000g. The supernatants were loaded onto a NiNTA column (Qiagen) at 4˚ C, which was subsequently washed with Ni Wash Buffer (50 mM Tris pH 8.0, 300 mM NaCl, 10 mM imidazole, 10% glycerol, 0.5 mM DDM) and bound proteins eluted with Ni Elution Buffer (50 mM Tris pH 8.0, 300 mM NaCl, 250 mM imidazole, 10% glycerol, 0.5 mM DDM). Eluate from the NiNTA column was concentrated before separation on a Superdex 200 gel filtration column (GE Healthcare) equilibrated in 20 mM Tris pH 8.0, 150 mM NaCl, 10% glycerol, and 0.5 mM DDM. SDS-PAGE analysis of the resulting sample revealed the presence of four proteins, which were confirmed by trypsin digestion and tandem mass spectrometry (MS/MS) to be MlaF, MlaE, MlaD and MlaB. For cryo-EM, the sample in DDM was exchanged into amphipol A8-35 prior to size exclusion chromatography. Briefly, protein at ∼1mg/mL was incubated for ∼5 hours with 3-fold excess by mass of A8-35. Detergent from the buffer was then absorbed by incubation with Bio-Beads (Bio-Rad) overnight at 15 mg Bio-Beads per mL of solution. The sample was filtered to remove Bio-Beads and subjected to size exclusion chromatography. The protein quality was then assessed by negative stain EM prior to cryo-EM grid preparation.

### Expression and purification of PqiB

DNA corresponding to mature, signal peptide-cleaved PqiB (residues 39-546) was amplified from the MG1655 genome and cloned by Gibson assembly (Gibson et al., 2009) into a custom pET vector resulting in an N-terminal 6xHis tag and TEV protease cleavage site. The resulting plasmid (pBE1163) was transformed into Rosetta 2 cells (Novagen). For expression, overnight cultures of Rosetta 2/pBE1163 were grown at 37˚ C with shaking to an OD600 of ∼0.9, then induced by addition of IPTG to a final concentration of 1 mM and continued incubation overnight shaking at 15˚ C. Cultures were harvested by centrifugation, and pellets were resuspended in lysis buffer (50 mM Tris pH 8.0, 300 mM NaCl, 10 mM imidazole). Cells were lysed by two passes through an Emulsiflex-C3 cell disruptor (Avestin), then centrifuged at 38,000g to pellet cell debris. The clarified lysates were incubated with NiNTA resin (Qiagen) at 4˚ C, which was subsequently washed with Ni Wash Buffer (50 mM Tris pH 8.0, 300 mM NaCl, 10 mM imidazole) and bound proteins eluted with Ni Elution Buffer (50 mM Tris pH 8.0, 300 mM NaCl, 250 mM imidazole). PqiB containing fractions eluted from the NiNTA column were pooled and concentrated before separation on a Superdex 200 gel filtration column (GE Healthcare) equilibrated in 20 mM Tris pH 8.0 and 150 mM NaCl.

### Expression and purification of YebT

DNA encoding the *yebST* operon was amplified from the MG1655 genome and cloned by Gibson assembly (Gibson et al., 2009) into a custom pET vector resulting in a bicistronic YebS-YebT co-expression construct with a C-terminal 6xHis tag on YebT. The resulting plasmid (pBE1287) was transformed into Rosetta 2 cells (Novagen). Expression and purification were carried out as described for PqiB above. Though this construct co-expresses YebS and YebT, only significant amounts of full-length YebT were obtained under our expression conditions.

## Electron microscopy (EM) data collection

### Negative stain EM data

To prepare grids for negative stain EM, sample after size exclusion chromatography was applied to a freshly glow discharged carbon coated 400 mesh copper grids and blotted off. Immediately after blotting, a 2% uranyl formate solution was applied for staining and blotted off. The stain was applied five times per sample. Samples were allowed to air dry before imaging. Data were collected on a Tecnai T12 microscope (FEI) equipped with a 4K x 4K CCD camera (UltraScan 4000, Gatan) at a nominal magnification of 52,000 corresponding to a pixel size of 2.21 Å2/pixel on the specimen, and a defocus range of 1 – 1.7 uM underfocus.

### Cryo EM of MlaFEDB

For cryo-EM, 3μL of MlaFEDB complex at a concentration of ∼1 mg/mL and containing amphipol A8-35 was applied to quantifoil holey carbon grids (1.2/1.3, 400 mesh). Grids were blotted for 6 s at 4 °C and plunge frozen in liquid ethane using a Vitrobot Mark III. Cryo-EM images were collected on a Titan Krios, FEI company (Janelia Research Campus, “Krios 2”) operated at 300 kV and equipped with a K2 Summit direct electron detector camera (Gatan). Images were recorded using super-resolution counting mode following an established protocol (Li et al., 2013). Specifically, images were recorded at a nominal magnification of 20,000 x, corresponding to a calibrated super-resolution pixel size of 1.02 Å on the specimen and 0.51 Å for super-resolution images. The dose rate on the camera was set to be › 8 counts (corresponding to › 9.9 electron) per physical pixel per second. The total exposure time was 6 s, leading to a total accumulated dose of 41 electrons per Å2 on the specimen. Each image was fractionated into 30 subframes, each with an accumulation time of 0.2 s per frame. All dose-fractionated cryo-EM images were recorded using SerialEM (Mastronarde, 2005). Images were recorded with a defocus in the range from 1.5 to 3.0 μm.

### Cryo EM of PqiB

For cryoEM, PqiB protein at a concentration of ∼0.8 mg/mL was used. Although this is a soluble construct, 0.1 mg/mL amphipol A8-35 was added prior to freezing the grids. 3ML of sample was applied to quantifoil holey carbon grids (1.2/1.3, 400 mesh). Grids were blotted for 6 s at 4 °C and plunge frozen in liquid ethane using a Vitrobot Mark III. Cryo-EM images were collected on a Titan Krios, FEI company (Janelia Research Campus, “Krios 2”) operated at 300 kV and equipped with a K2 Summit direct electron detector camera (Gatan). Data collection was carried out using SerialEM. Images were recorded at a nominal magnification of 22,500 x, corresponding to a super-resolution pixel size of 1.31 Å on the specimen and 0.655 Å for super-resolution images. The rate on the camera was set at 10 electrons per pixel per second. Each image was fractionated into 50 frames with an exposure of 280 ms/frame, resulting in a total exposure time of 14 seconds and a total dose of ∼80 electrons /Å2. A total of ∼1700 images were recorded during an ∼2-day period with a defocus range of 0.8 to 2.0 μm.

## EM Data processing

### Negative stain EM data

Images were binned, resulting in a pixel size of 4.42 Å2/pixel. Particles were picked automatically using template-based picking. Simplified Application Managing Utilities for EM Labs (SAMUEL, http://liao.hms.harvard.edu/samuel) scripts were used for image preprocessing, particle picking and 2D classification, as described by Ru *et al* in Supplemental Experimental Procedures (Ru et al., 2015).

After a satisfactory stack of particles were obtained, an initial model was obtained using the software simple 2.0 (Elmlund and Elmlund, 2012; Elmlund et al., 2013). 3D classification was performed on the stack in RELION (Scheres, 2012a, 2012b; Scheres and Chen, 2012) using the initial model generated from simple 2.0 filtered to 50 Å as an initial model. The data were divided into 4 classes, and each resulting class was refined using the auto-refine procedure in RELION.

### Cryo EM processing of MlaFEDB

Images were binned, resulting in a pixel size of 3.03 Å2/pixel for cryo-EM data of the MlaFEDB complex. Particles were picked automatically using template-based picking. Simplified Application Managing Utilities for EM Labs (SAMUEL, http://liao.hms.harvard.edu/samuel) scripts were used for image preprocessing, particle picking and 2D classification, as described by Ru *et al* in Supplemental Experimental Procedures (Ru et al., 2015). After a satisfactory stack of particles were obtained, 2D classification was also carried out in RELION. The resulting stack of particles was subjected to iteratve 3D classification and the auto-refine procedure in RELION until the most homogeneous set of particles was obtained. Some compositional heterogeneity was present, which resulted in a large amount of top views of MlaD alone (not in complex with MlaE, F and B). These particles were separated out early in procedure. The remaining particles of the MlaFEDB complex also contain conformational heterogeneity, and possible compositional heterogeneity with respect to the bound nucleotide, as the nucleotide state was not controlled in this dataset. No exogenous ATP, ADP or ATP analog was added, so we expect the sample to be largely in the “apo” state. However, it is possible that endogenous nucleotide may be bound in some particles. Regardless, the orientation between the permease subunit (MlaE), the ATP-binding domains (MlaF) and the potential regulatory subunits (MlaB) shows some flexibility. Therefore, while several 2D classes show detailed features, these are lost in the 3D reconstruction, even after 3D classification. Likely a much larger dataset with a more homogenous sample will be required to reach higher resolution and identify the pseudo-atomic details of the complex.

### Cryo EM processing of PqiB

Drift correction was carried out using MotionCor (Zheng et al., 2016), as MotionCor2 was still under development at this time. CTF estimation was performed using the standalone version of Gctf (Zhang, 2016). Particles were picked using Gautomatch (currently unpublished; developed by Zhang K, MRC Laboratory of Molecular Biology, Cambridge, UK, http://www.mrc-lmb.cam.ac.uk/kzhang/Gautomatch/). Briefly, a few hundred particles were picked from ∼5 micrographs. Particles were extracted in Relion and 2D classificatoin in Relion (Scheres, 2012a, 2012b; Scheres and Chen, 2012) was used for alignment and classification to generate templates for further particle picking. Template-based picking was then carried out on all micrographs using Gautomatch. 2D classification in Relion was used to choose ∼357,000 “good” particles, and iterative 3D classification and 3D autorefine were used to generate the final reconstruction. The particles were refined with C6 symmetry as well as no symmetry, which resulted in similar reconstructions, but higher quality maps with imposition of C6 symmetry. The protein can roughly be divided into 2 parts: the “barrel” and the “needle” (as described in the main text. As is evident from the 2D classes, the barrel is fairly stable, and tends to drive the alignment, whereas the needle is flexible relative to the barrel. To obtain a better reconstruction for the needle, the box size was increased and particles were re-extracted to include the whole needle, and a mask was applied, either to Ring3 and the needle, or the needle only during refinement. These reconstructions provided additoinal detail for the needle part of PqiB. Post-processing was done in Relion, and using the automask and autoB-factor options usually resulted in the highest quality map.

### Model building and refinement for PqiB

3 copies of the hexameric MlaD ring were docked into the EM density map and fit as rigid bodies in Chimera (Pettersen et al., 2004). Individual domains were then fit as rigid bodies in Coot (Emsley et al., 2010). Regions of poor agreement between the model and map were pruned and then refined using phenix.real_space_refine (Adams et al., 2010). The PqiB backbone model was rebuilt and the sequence register assigned by manual inspection of the EM density map in Coot (Emsley et al., 2010), along sequence and structural alignments to MlaD. Additional density for the C-terminal helical region was fit manually using idealized α-helices. The final model was refined using phenix.real_space_refine (Adams et al., 2010) against 2 different EM density maps: one masked over the entire model, which provided the best resolution of the PqiB barrel, and a second map masked on the MCE3 ring and needle, which yielded the highest quality maps for the needle. Geometry of the final model was validated using MolProbity (Chen et al., 2010).

<u>Needle</u>. While the positions and overall twist of the C-terminal helices was readily apparent in our maps, the lower local resolution in this area precluded the assignment of the sequence register within the needle. Consequently, these regions have only been modeled as poly-alanine,

<u>18-stranded β-barrel lining interior of PqiB barrel</u>. Considering only the packing of the core MCE domains in PqiB, large holes between subunits would allow for solvent and some small molecules to diffuse in and out of the PqiB barrel. However, in the regions where these holes exist (between the MCE1 and MCE2 rings; and between the MCE2 and MCE3 rings), our EM maps reveal that the interior of the barrel is lined with 18 tubes of density that run roughly parallel to the long/six-fold axis of PqiB. These densities exhibit the characteristic “pleated” shape of β-strands, with spacing between tubes of density consistent with a curved β-sheet, suggesting that the 18 densities correspond to an 18-stranded β-barrel that seals the gaps between the MCE rings. Consistent with this, densities resembling side chains protrude off from the strands, alternating between the lumenal and exterior faces of the barrel. In both parts of the structure lined by these 18 densities (between the MCE1 and MCE2 rings; and between the MCE2 and MCE3 rings), twelve of the eighteen β-strands, two from each of the 6 subunits in the ring, can be modeled as a β-hairpin emanating from the β5-6 loops in the MCE1 and MCE2 domains of PqiB, which are much longer than the corresponding β5-6 loop of MlaD of the MCE3 domain of PqiB. In the remaining 6 β-strand densities lining the pore, we have built polyalanine models, as we are currently unable to determine where these residues lie within the primary sequence, or if they may be derived from another polypeptide co-purifying with PqiB. Indeed, 3D classification applied during the reconstruction of PqiB yielded two major classes that differed primarily in the presence or absence of strong density for this 18-stranded β-barrel, and it is possible that the 3D class with poor density in this region is missing an a polypeptide the contributes the remaining 6 strands and thereby stabilizes the pore lining. Future work and likely higher resolution structures will be needed to resolve this ambiguity.

### Identification of inter-subunit restraints in MlaFEDB complex using co-evolving residue pairs

A multiple sequence alignment was generate using HHblits (Remmert et al., 2012) with UniProt database (UniProt Consortium, 2014) from 2015_06 for each of the Mla components. To prevent paralogous copies of each genes from corrupting the final alignments, stringent e-values were used. For MlaE, MlaF and MlaD evalues of 1E-20, 1E-40 and 1E-20 were used respectfully. The joint alignment was restricted to genes found within the same operon (maximum of two genes apart in the same contig). The genomic distance was determined by mapping the UniProt IDs back to their respective contiguous sequences from the ENA database (Leinonen et al., 2010). The alignments were then filtered to remove sequences that don't cover at least 75 % of the query, positions that have more than 50% gaps and redundancy was reduced to 90%. GREMLIN was then run with default parameters to predict the top inter-co-evolving residue pairs (Ovchinnikov et al., 2014). Residue pairs with predicted probability of 80% or higher are shown in Figures S3C and S3D mapped on the model structures.

The previously predicted structure of MlaE dimer (Ovchinnikov et al., 2015) was refined using the rosetta hybrid protocol (Song et al., 2013) in context of MlaF dimer and co-evolution restraints. The MlaF dimer was modeled using outward-facing Maltose/maltodextrin import ATP-binding protein MalK conformation [3RLF; (Oldham and Chen, 2011)].

### Expression and purification of MlaC

DNA corresponding to mature, signal peptide-cleaved MlaC (residues 22-211) was amplified from the MG1655 genome and cloned by Gibson assembly (Gibson et al., 2009) into a custom pET vector resulting in an N-terminal 6xHis tag and TEV protease cleavage site. The resulting plasmid (pBE1203) was transformed into Rosetta 2 cells (Novagen). For expression, overnight cultures of Rosetta 2/pBE1203 were grown at 37˚ C with shaking to an OD600 of ∼0.9, then induced by addition of IPTG to a final concentration of 1 mM and continued incubation overnight shaking at 15˚ C. Cultures were harvested by centrifugation, and pellets were resuspended in lysis buffer (50 mM Tris pH 8.0, 300 mM NaCl, 10 mM imidazole). Cells were lysed by two passes through an Emulsiflex-C3 cell disruptor (Avestin), then centrifuged at 38,000g to pellet cell debris. The clarified lysates were incubated with NiNTA resin (Qiagen) at 4˚ C, which was subsequently washed with Ni Wash Buffer (50 mM Tris pH 8.0, 300 mM NaCl, 10 mM imidazole) and bound proteins eluted with Ni Elution Buffer (50 mM Tris pH 8.0, 300 mM NaCl, 250 mM imidazole). MlaC containing fractions eluted from the NiNTA column were pooled and concentrated before separation on a Superdex 200 gel filtration column (GE Healthcare) equilibrated in 20 mM Tris pH 8.0 and 150 mM NaCl.

### Crystallization and structure determination of MlaC

Gel filtration fractions containing purified MlaC proteins were concentrated to 10-60 mg/mL and sitting drop, vapor-diffusion crystallization trials were conducted using the JCSG Core I-IV screens (Qiagen). Crystals of MlaC grew from drops consisting of 100 nL protein plus 100 nL of a reservoir solution consisting of 0.1 M citric acid pH 3.5 and 1.6 M ammonium sulfate, and were cryoprotected by supplementing the reservoir solution with 37.5% ethylene glycol. Native diffraction data was collected at ALS beamline 8.3.1, and indexed to P212121 and reduced using XDS (Table S1) (Kabsch, 2010). The structure was phased by molecular replacement using Phaser (McCoy et al., 2007). Initial attempts to use the crystal structures of MlaC homologs as search models (PDB IDs: 2QGU and 4FCZ) were unsuccessful. However, we were able to obtain a few clear solutions using the following approach. We created a homology model of MlaC with Phyre2 (Kelley et al., 2015), based on 2QGU as a template. This homology model rebuilt using MR-Rosetta (DiMaio et al., 2011), and the resulting 1000 rebuilt models were tested as search models using Phaser (McCoy et al., 2007). A small number of these models gave clear solutions, which were adjusted in Coot (Emsley et al., 2010) and refined using Phenix (Afonine et al., 2012). In the later stages of refinement, two phosphatidic acid molecules were built into clearly defined electron density in the core of each of the two MlaC monomers in the asymmetric unit.

### Expression and purification of MlaA-OmpF complex

DNA corresponding to mature, signal peptide-cleaved MlaA (residues 1-251) was amplified from the MG1655 genome and cloned by Gibson assembly (Gibson et al., 2009) into a custom pET vector resulting in an C-terminal 6xHis tag. The resulting plasmid (pBE1242) was transformed into Rosetta 2 cells (Novagen). For expression, overnight cultures of Rosetta 2/pBE1242 were grown at 37˚ C with shaking to an OD600 of ∼0.9, then induced by addition of IPTG to a final concentration of 1 mM and continued incubation for ∼4hr shaking at 15˚ C. Cultures were harvested by centrifugation, and pellets were resuspended in lysis buffer (50 mM Tris pH 8.0, 300 mM NaCl, 10 mM imidazole, 10% glycerol). Cells were lysed by two passes through an Emulsiflex-C3 cell disruptor (Avestin), then centrifuged at 38,000g to pellet cell debris. The clarified lysates transferred to ultracentrifuge tubes and centrifuged at 200,000g to pellet membrane fraction. Membrane pellets were resuspended in lysis buffer by sonication, then n-Dodecyl-β-D-Maltopyranoside (DDM; Anatrace) was added to a final concentration of 25 mM. After incubation for 1hr with rocking at 4˚ C, the DDM solublized membrane fraction was centrifuged at 200,000g. The supernatants were loaded onto a NiNTA column (Qiagen) at 4˚ C, which was subsequently washed with Ni Wash Buffer (50 mM Tris pH 8.0, 300 mM NaCl, 10 mM imidazole, 10% glycerol, 0.5 mM DDM) and bound proteins eluted with Ni Elution Buffer (50 mM Tris pH 8.0, 300 mM NaCl, 250 mM imidazole, 10% glycerol, 0.5 mM DDM). MlaA containing fractions eluted from the NiNTA column were pooled and concentrated before separation on a Superdex 200 gel filtration column (GE Healthcare) equilibrated in 20 mM Tris pH 8.0, 150 mM NaCl, 10% glycerol, and 0.5 mM DDM. SDS-PAGE analysis of the resulting sample revealed the presence of two proteins in roughly 1:1 stoichometry, which were confirmed by trypsin digestion and tandem mass spectrometry (MS/MS) to be MlaA and OmpF.

### Protein-protein interactions by biolayer interferometry

DNA corresponding to mature, signal peptide-cleaved MlaC (residues 22-211) was amplified from the MG1655 genome and cloned by Gibson assembly (Gibson et al., 2009) into a custom pET vector resulting in an N-terminal 6xHis tag, biotinylation site (amino acid sequence: GGGLNDIFEAQKIEWHE), and TEV protease cleavage site. The resulting plasmid (pBE1230) was transformed into Rosetta 2 cells (Novagen), and tagged MlaC was expressed and purified essentially as described above. Purified MlaC at ∼2 mg/mL was biotinylated by the addition of 25μg BirA enzyme/mg total protein, purified essentially as previous described (Ekiert et al., 2011). The reactions were carried out in the following buffer: 100mM Tris pH 8.0, 10mM ATP, 10mM MgOAc, 50μM biotin. The biotinylation reactions were incubated at 4 ˚C overnight. Biotinylated MlaC was purified by size exclusion chromatography, and concentrated to ∼5-20 mg/mL.

Biotinylated MlaC was diluted to 50 μg/mL in 1x kinetics buffer (1x PBS, pH 7.4, 0.01% BSA, and 0.002% Tween 20) and loaded onto streptavidin coated biosensors (ForteBio) and incubated with varying concentrations of MlaFEDB complex, MlaD, or MlaA-OmpF. All binding data were collected on an Octet RED384 instrument at 25 ˚C. In each case, an empty sensor (no MlaC loaded) was included as a negative control to test for non-specific interactions of MlaFEDB complex, MlaD, and MlaA-OmpF.

**Figure S1.**
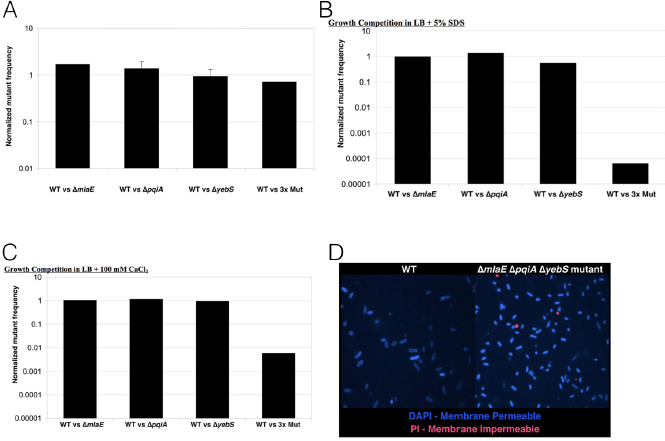
Comparison of WT and MCE mutant *E. coli* strain growth and OM barrier phenotypes, related to Figure 1. (A) After mixing equal volumes of WT and MCE mutant overnight cultures, mixed cultures were diluted 1:1000 in fresh LB and incubated ∼16hr at 37˚ C with shaking. The fraction of mutant cells (mutant cells divided by total cells) was estimated before and after growth competition by plating dilutions on LB (total cells) and LB+Kan (mutant cells). The ratios presented here represent the change in the fraction of mutant cells between the initial mixed culture and the post-competition culture [(fraction mutant T16hr) / (fraction mutant T0)]. Values above 1 indicate a growth advantage in the mutant, while values below 1 indicate a growth defect in the mutant. For all the mutants, the frequency of mutant cells changed by less than 2-fold between the input and post-competition cultures, suggesting MCE genes are dispensable for in vitro growth in rich media. (B and C) After mixing equal volumes of WT and MCE mutant overnight cultures, mixed cultures were diluted 1:1000 in fresh LB+5% SDS (B) or LB+100 mM CaCl2 (C), incubated ∼16 hr at 37C with shaking, and analyzed as described above. Addition of 5% SDS or 100 mM CaCl2 had little effect on the growth of any of the single mutants, but dramatically reduces the frequency of the triple mutant. (D) Live cells growing exponentially in LB were co-stained with DAPI and PI for 30 minutes then imaged. The culture of the triple mutant had more PI-positive cells than the wild-type, though the vast majority of cells in both cultures were PI-negative, indicating that the inner membrane remains intact. DAPI staining of the triple mutant was brighter than the wild-type, with clear nucleoid localization, while the staining of wild-type cells under these conditions was strongest at the cell periphery (cell wall/membrane staining). Consistent with the role of the OM as a barrier to the entry of small molecules into the cell, DAPI entry into WT cells appears to be inefficient, while in the triple mutant the OM is compromised, allowing efficient nucleoid staining.

**Figure S2.**
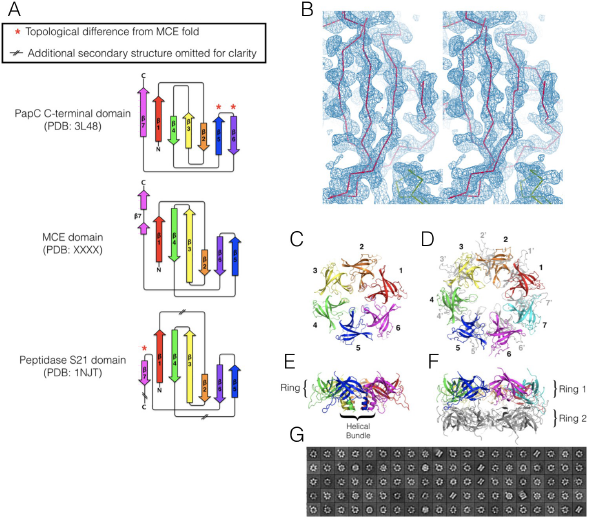
Structure of MCE protein, MlaD, related to Figure 2. (A) Topology diagrams highlighting the differences between the MCE domain and the closest structural neighbors in the PDB. Adapted from Pro-Origami diagrams (Stivala et al., 2011)). (B) Initial electron density map for the core MCE domain of MlaD [32-140] after phasing and density modification using SHELX C/D/E pipeline. Map is contoured at 2σ and displayed as a blue mesh. The Cα trace of the final model is shown in red for comparison. (C), the full-length MlaD ectodomain, which includes an additional ∼45 amino acids at the C-terminus that associate to form a helical bundle, forms a single hexameric ring. (D), in contrast, the core MCE domain forms a tetradecamer, consisting of two rings of seven MCE domains stacked on top of one another with their pores roughly aligned (subunits of upper ring colored, bottom ring gray). (E), 90˚ rotated view of the full-length, hexameric MlaD ectodomain. (F), 90˚ rotated view of the tetradecameric MlaD core MCE domain, colored as in (B). (G), 2D class averages from single particle, negative stain electron microscopy are also consistent with a 14 subunit, stacked ring organization for the MCE core domain from MlaD, suggesting that the 14-mer observed in the crystal is also stable in solution. We believe the hexamer is the predominant and perhaps the only functional form *in vivo*, for three reasons. 1) The complete periplasmic domain crystal structure is hexameric, and it is also hexameric in the context of the larger MlaFEDB complex. Second, PqiB and YebT, two other MCE proteins from *E. coli*, are only hexameric. Third, the “head-to-head” arrangement of the two stacked rings in the 14-mer places the C-terminal helical bundle from each ring at the interface, resulting in major steric overlap.

**Figure S3.**
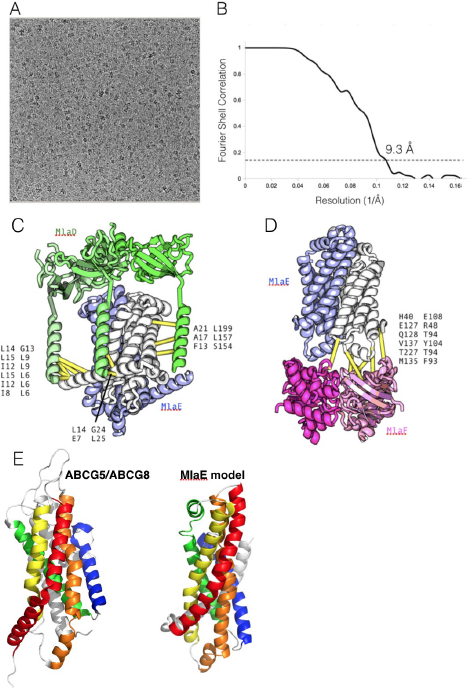
Cryo EM and computational modeling of the MlaFEDB complex, related to Figure 3. (A) Representative raw micrograph of MlaFEDB using a K2 camera. (B) Gold standard FSC curve of final reconstruction. Horizontal line indicates 0.143 FSC. (C and D) Location of strongly co-evolving residue pairs in the MlaFEDB complex. (C) Co-evolving residue pairs suggestive of potential contacts between MlaD (left column) and MlaE (right column). (D), Co-evolving residue pairs suggestive of potential contacts between MlaE (left column) and MlaF column). (E) Comparison of the *de novo* model of MlaE with the crystal structure of the permease domain of the ABCG5/ABCG8 transporter structure (PDB code 5DO7). Helices are colored from red to blue from the N-terminus to the C-terminus, indicating the similar fold and topology between these two proteins. An additional N-terminal helix in MlaE that is not found in ABCG5/ABCG8 and additional C-terminal helices in ABCG5/ABCG8 that are not present in MlaE are colored light grey.

**Figure S4.**
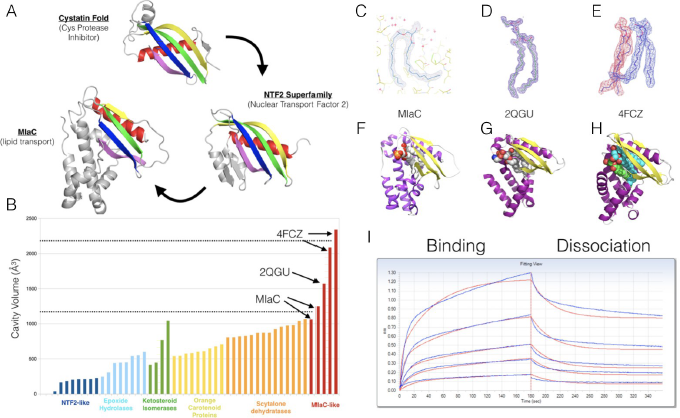
Structural analyses and binding data for MlaC, related to Figure 4. (A), Comparison of MlaC with cystatin and other NTF2 superfamily members. In contrast to cystatins, where a single α-helix (red) packs tightly against one face of the β-sheet, the NTF2 superfamily extends the sheet with 2 additional β-strands while adding two α-helices, creating a hydrophobic pocket between the sheet and helices. MlaC adds several addition helices to this core cystatin/NTF2 fold. (B), The hydrophobic pocket varies in volume across the diverse members of the NTF2 superfamily, from a modest depression in NTF2 itself that mediates interaction with Ran-GDP (dark blue bars), to a prominent active site cleft in many enzymes where it can accommodate small hydrophobic substrates, such as steroids in the ketosteroid isomerases (green bars). Compared with other members of the NTF2 superfamily, MlaC and the related 2QGU and 4FCZ protein structures have some of the largest cavities observed to date (red bars). (C-H) All three MlaC-like structures determined to date have bound lipid. 2Fo-Fc electron density map contoured at 1 sigma for MlaC (C), 2QGU (D), and 4FCZ (E), showing only the density corresponding to the bound lipids. In (E), the density was segmented and colored red and blue to indicate the regions corresponding to the two putative diacylglycerol-like groups. The bound lipids are similarly oriented in each structure [white spheres for MlaC (F) and 2QGU (G); and green and cyan spheres for the two diacylglycerol-like groups of 4FCZ (H)]. (I) MlaD interacts directly with biotinylated MlaC protein immobilized on a streptavidin-coated biosensor over a range of MlaD concentrations, as assayed by biolayer interferometry.

**Figure S5.**
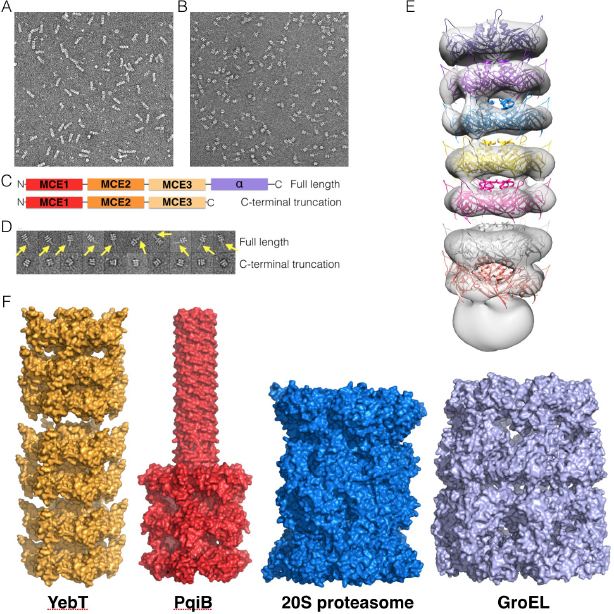
Negative-stain single particle EM of YebT and PqiB, related to Figure 5. (A and B) Representative raw micrograph of negative-stained YebT (A) and PqiB (B). (C) Schematic of constructs that were used. Top, full length construct of PqiB periplasmic domain; Bottom, construct in which the C-terminal domain, predicted to be helical, has been truncated. (D) Representative single particles from negative stain EM data for the constructs shown in (C). Yellow arrows point to the density corresponding to a needle-like projection. This density is missing from the C-terminal truncated construct, suggesting that the needle-like projection is formed by the C-terminal domain. (E) Negative stain single particle EM density map for full-length YebT (semi-transparent white surface). The small, roughly spherical density at bottom likely corresponds to the N-terminal transmembrane helices from each subunit, which would be anchored in the inner membrane. Above this, 7 stacked MCE rings (colored ribbons) would extend into the periplasmic space, towards the outer membrane. (F) Comparison of YebT and PqiB EM models with other supramolecular barrels: 20S proteasome and GroEL. All models are depicted to scale. The the barrels of YebT and PqiB are narrower than the proteasome and GroEL, but YebT's is considerably longer.

**Figure S6.**
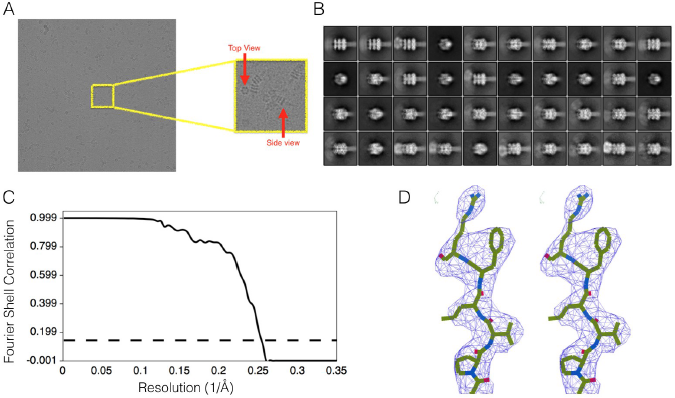
Cryo EM reconstruction of PqiB at 3.96 Å resolution, related to Figure 7. (A) Representative raw micrograph of PqiB using a K2 camera. (B) Representative 2D class averages for PqiB. (C) Gold standard FSC curve of final reconstruction. Horizontal line indicates 0.143 FSC. (D) Stereo view of a representative region of the PqiB EM density map (blue mesh) and final atomic model for residues Pro175-Val176-Leu177-Phe178-Arg179.

**Figure S7.**
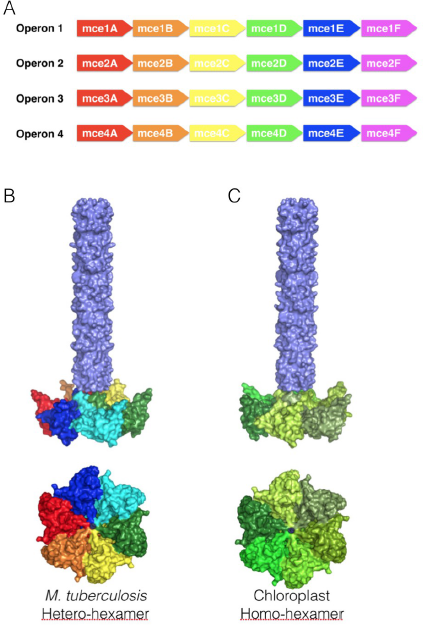
Possible architectures of MCE domains proteins from *M. tuberculosis* and chloroplasts, related to Figure 7. (A) The *M. tuberculosis* genome has 4 MCE operons, each encoding 6 MCE genes. Based upon the hexameric assemblies formed by *E. coli* MCE proteins, we propose that each MCE operon in *M. tuberculosis* MCE codes for a single, hetero-hexameric assembly, with 6 unique chains---one from each MCE gene. (B) MCEs from *M. tuberculosis* encode a single MCE domain followed by a long C-terminal helical region. Based on the similar architecture of PqiB's final MCE3 domain and helical C-terminal domain, we predict that *M. tuberculosis* MCE proteins will form a single hetero-hexameric ring with a long needle-like projection. (C) The chloroplasts of plants and other photosynthetic eukaryotes contain a single MCE gene encoding a protein with a single MCE domain followed by a long C-terminal helical region (called TGD2 in *A. thaliana*). We predict that TGD2 will form a single homo-hexameric ring with a long needle-like projection.

**Table S1.** Crystallographic data collection and refinement statistics.

|  | MlaD<br>[32-140; MCE core domain] | MlaD<br>[32-183; periplasmic domain] | MlaC |
| --- | --- | --- | --- |
| **Data collection** |  |  |  |
| Space group | C2 | C2 | P2<sub>1</sub>2<sub>1</sub>2<sub>1</sub> |
| Cell dimensions |  |  |  |
| *a*, *b*, *c* (Å) | 139.7, 103.8, 77.2 | 122.4, 63.8, 91.2 | 33.6, 115.9, 133.2 |
| $\alpha$, $\beta$, $\gamma$ (°) | 90.0, 111.0, 90.0 | 90.0, 130.0, 90.0 | 90.0, 90.0, 90.0 |
| Wavelength (Å) | 1.116 | 1.116 | 1.116 |
| Resolution (Å) | 50-2.15 (2.21-2.15)<sup>1</sup> | 50-2.85 (2.92-2.85)<sup>1</sup> | 50-1.50 (1.54-1.50)<sup>1</sup> |
| Observations | 421,073 | 46,094 | 599,263 |
| Unique reflections | 55,590 | 12,551 | 83,295 |
| Redundancy | 7.6 (7.6) | 3.7 (3.7) | 7.2 (7.1) |
| Completeness (%) | 99.3 (98.6) | 97.8 (98.9) | 98.5 (96.1) |
| CC*<sup>2</sup> | 1.00 (0.80) | 1.00 (0.79) | 1.00 (0.85) |
| CC1/2<sup>2</sup> | 0.99 (0.46) | 1.00 (0.47) | 1.00 (0.57) |
| *I* / $\sigma I$ | 15.4 (0.9) | 14.9 (1.0) | 22.2 (1.5) |
| *R*<sub>meas</sub> | 0.09 (2.46) | 0.05 (1.76) | 0.05 (1.40) |
| *R*<sub>pim</sub> | 0.04 (0.88) | 0.03 (0.91) | 0.02 (0.51) |
| **Refinement** |  |  |  |
| Resolution (Å) | 50-2.15 | 50-2.85 | 50-1.50 |
| Reflections (work) | 48,118 | 11,005 | 77156 |
| Reflections (free) | 1810 | 589 | 1,894 |
| *R*<sub>work</sub> (%) / *R*<sub>free</sub> (%) | 20.7 / 24.4 | 23.3 / 28.8 | 16.9 / 18.5 |
| No. atoms |  |  |  |
| Protein | 5,552 | 2,513 | 3,147 |
| Ligand/ion | 0 | 6 | 80 |
| Water | 126 | 0 | 264 |
| *B*-factors |  |  |  |
| Protein | 70.3 | 124.7 | 29.9 |
| Ligand/ion | na | 170.8 | 52.4 |
| Water | 53.6 | na | 39.3 |
| R.m.s. deviations |  |  |  |
| Bond lengths (Å) | 0.003 | 0.003 | 0.009 |
| Bond angles (°) | 0.94 | 0.79 | 1.23 |
| Ramachandran |  |  |  |
| Favored | 97.8 | 97.5 | 99.0 |
| Outliers | 0.7 | 0.3 | 0.0 |
| **PDB Code**<sup>3</sup> | aaaa<sup>3</sup> | bbbb<sup>3</sup> | cccc<sup>3</sup> |

<sup>1</sup> Values in parentheses are for highest-resolution shell.<sup>2</sup> See (Karplus and Diederichs, 2012).<sup>3</sup> Coordinates and structure factors will be deposited in the Protein Databank and released upon publication.

**Table S2.** Crystallographic data collection and MAD phasing statistics for MlaD structure.

| MlaD [32-140; MCE core domain] |  |  |
| --- | --- | --- |
| **Data collection** |  |  |
| Space group | C2 |  |
| Cell dimensions |  |  |
| *a*, *b*, *c* (Å) | 139.3, 103.6, 77.6 |  |
| $\alpha$, $\beta$, $\gamma$ (°) | 90.0, 111.1, 90.0 | |
|  | Peak | Remote |
| Wavelength (Å) | 0.980 | 0.957 |
| Resolution (Å) | 50-2.80 (2.87-2.80)<sup>1</sup> | 50-2.80 (2.87-2.80)<sup>1</sup> |
| Observations | 962,235 | 962,435 |
| Unique reflections | 49,671 | 49,681 |
| Redundancy | 19.4 (19.5) | 19.4 (19.5) |
| Completeness (%) | 99.6 (99.9) | 99.6 (99.9) |
| CC1/2<sup>2</sup> | 1.00 (0.67) | 1.00 (0.72) |
| *I* / $\sigma I$ | 11.9 (1.6) | 11.8 (1.7) |
| *R*<sub>meas</sub> | 0.32 (2.39) | 0.30 (2.25) |
| FOM | 0.55 |  |

<sup>1</sup> Values in parentheses are for highest-resolution shell.
<sup>2</sup> See (Karplus and Diederichs, 2012).

**Table S3.**
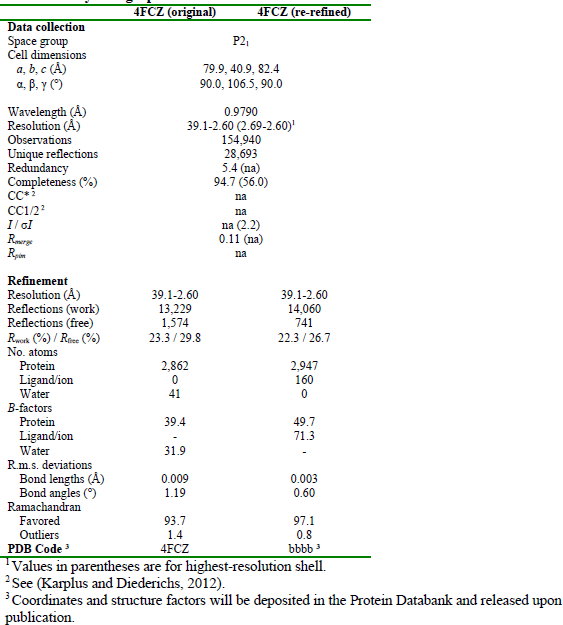
**Crystallographic data collection and refinement statistics for re-refined 4FCZ.**

**Table S4.**
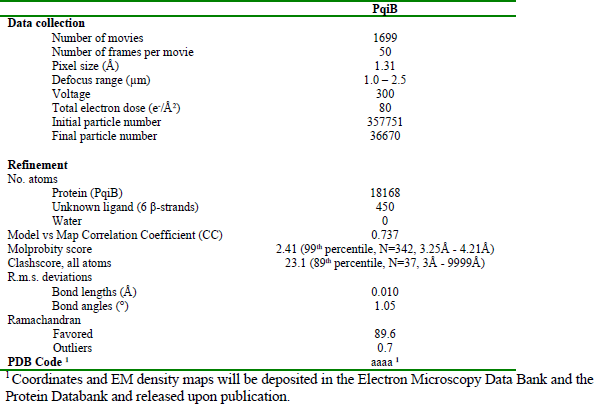
**Cryo EM data collection and refinement statistics for PqiB.**

